# Enduring consequences of perinatal fentanyl exposure in mice

**DOI:** 10.1101/735464

**Authors:** Jason B. Alipio, Adam T. Brockett, Megan E. Fox, Stephen S. Tennyson, Coreylyn A. deBettencourt, Dina El-Metwally, Nikolas A. Francis, Patrick O. Kanold, Mary Kay Lobo, Matthew R. Roesch, Asaf Keller

## Abstract

Opioid use by pregnant women is an understudied consequence associated with the opioid epidemic, resulting in a rise in the incidence of neonatal opioid withdrawal syndrome (NOWS), and lifelong neurobehavioral deficits that result from perinatal opioid exposure. There are few preclinical models that accurately recapitulate human perinatal drug exposure, and few focus on fentanyl, a potent synthetic opioid that is a leading driver of the opioid epidemic. To investigate the consequences of perinatal opioid exposure, we administered fentanyl to mouse dams in their drinking water throughout gestation and until litters were weaned at postnatal day (PD) 21. Fentanyl-exposed dams delivered smaller litters, and had higher litter mortality rates compared to controls. Metrics of maternal care behavior were not affected by the treatment, nor were there differences in dams’ weight or liquid consumption throughout gestation and 21 days postpartum. Twenty-four hours after weaning and drug cessation, perinatal fentanyl-exposed mice exhibited signs of spontaneous somatic withdrawal behavior, and sex-specific weight fluctuations that normalized in adulthood. At adolescence (PD 35) they displayed elevated anxiety-like behaviors and decreased grooming, assayed in the elevated plus maze and sucrose splash tests. Finally, by adulthood (PD 55) they displayed impaired performance in a two-tone auditory discrimination task. Collectively, our findings suggest that perinatal fentanyl exposed mice exhibit somatic withdrawal behavior and changes across the lifespan reminiscent of humans born with NOWS.

## Introduction

The NIH has deemed opioid misuse a national health crisis, with a 30% increase in the rate of opioid overdoses since 2016.^1^ The Center for Disease Control and Prevention estimates the total economic burden of opioid misuse in the United States at $78.5 billion annually,^2^ underscoring the enormous impact opioid misuse has on our health and financial well-being. The overwhelming and rapid nature with which this crisis has materialized is concerning and further warrants a careful understanding of the health consequences of opioid misuse.

Within this growing crisis, individuals between the age of 18 and 25 have exhibited the largest increase in illicit opioid use (Substance Abuse and Mental Health Services Administration, 2013). It is concerning that women of reproductive age show increased incidence of use, given that opioids – both natural and synthetic – can readily cross the placenta and the blood-brain barrier. Indeed, *in utero* opioid exposure is associated with deleterious effects in developing offspring, with more dramatic effects on neurological function observed in infants relative to adults.^3^

There has been an exponential increase in the distribution of the synthetic opioid, fentanyl, which is routinely incorporated into frequently used narcotics, such as heroin.^4^ Fentanyl is 50 to 100 times more potent than morphine and is chiefly responsible for the recent increase in synthetic opioid-related overdose deaths.^5^ The effects that synthetic opioids have on the developmental trajectory of offspring have been incompletely studied, and are critically needed to develop effective treatment and prevention plans.

Women who used opioids during pregnancy have an increased rate of undergoing premature birth, stillbirth, and infant death.^6^ Withdrawal symptoms occur in 30-80% of neonates exposed to opioids *in utero*,^7^ and they exhibit decreased birth weight and body size.^8^ Consequences may be long lasting, as children and adolescents exposed to pre and peri-natal opioids have disruptions in stress reactivity, altered glucocorticoid levels, hyperactivity, impulsivity, and aggression.^9^ Some of these findings in humans have been corroborated and expanded upon by rodent models, that have also revealed brain abnormalities associated with opioid exposure. For example, prenatally-exposed rodents show abnormalities in motor coordination,^10^ anxiety,^11^increased depression-like symptoms, ^12^learning and memory impairments,^13^ altered reproductive function,^14^ drug sensitivity, and analgesic responses.^15^

Animal models offer distinct benefits for studying the consequences of perinatal opioid exposure. However, previous models have primarily focused on the effects of prenatal morphine exposure, and only during part of the gestational period that is developmentally similar to humans in the second trimester, specifically during the rats gestation days 11 to 18.^14,16–18^ Only a small number of studies model opioid exposure in humans throughout pregnancy by exposing pregnant dams during both gestation and weaning.^19–22^ Others have used continuous exposure to opioids, by way of implantable osmotic minipumps,^23^ a model that does not mimic intermittent use in humans. Few have exposed rodents to prenatal opioids other than morphine.^23–29^ Here, we describe a novel model of perinatal *fentanyl* exposure by administering fentanyl in the drinking water of pregnant mouse dams until litters are weaned at postnatal day 21. This protocol resembles the entire human gestational period^30^ and demonstrates its face validity to the human condition, in that it results in postnatal, adolescent and adult phenotypes reminiscent of those in exposed humans.

## Materials and methods

### Animals

All procedures were conducted in accordance with the *Guide for the Care and Use of Laboratory Animals* and approved by the Institutional Animal Care and Use Committees at the University of Maryland School of Medicine and College Park. Unless otherwise indicated, both male and female C57BL/6J mice were used, and bred in our facility. For auditory discrimination tasks we used the offspring of male CBA/C57BL/6J crossed with female Thy1-GCaMP6f/C57BL/6J mice since CBA strains do not exhibit age-related auditory deficits due to peripheral hearing loss,^31^ and were readily available from other ongoing calcium imaging studies. Separate cohorts of animals were used for each behavioral test to prevent possible crossover effects between tests. When copulatory plugs were identified, we removed the sires, and added fentanyl to the water hydration pouches (see below). Vehicle controls received water with 2% saccharin. Pouches were replenished weekly until litters were weaned at postnatal day (PD) 21. Offspring were housed 2 to 5 per cage, single-sex groups, in a temperature-and humidity-controlled vivarium. Food and water were available *ad libitum*, and lights were maintained on a 12-hour cycle.

### Fentanyl citrate

We used 10 μg/mL fentanyl citrate (calculated as free base) in 2% (w/v) saccharin, or 2% saccharin (vehicle control) in the drinking water. This concentration was selected, since it is the optimal concentration mice will readily self-administer without resulting in motor deficits,^32,33^ which is well below the mouse oral LD50 of 368 mg/kg (MSDS Cayman Chemical). C57BL/6 mice have an average daily water intake of 4 to 6 mL per day,^34^ which increases 3 to 4 times during pregnancy and lactation.^35^

### Maternal care behavior

We assessed maternal behavior as previously described.^36,37^ We carried out observations in the vivarium from PD 1 to 7, at daily sessions, between 9:00 and 10:30 AM, and between 2:00 and 3:30 PM. We scan sampled each dam in real time, once a minute for 30 minutes, yielding 30 scans per session; 2 sessions per day for 7 days yielded a total of 420 scans per dam. We used ethological parameters (Table 1): In nest behaviors comprised of: “licking/grooming”, “active nursing”, “passive nursing”, and “nest building” as manifestations of caring behavior; “self-grooming” as a manifestation of dam self-maintenance; and “pups out of nest” as a neglecting behavior. Out of nest behaviors were “eating/drinking” and “self-grooming” out of nest as self-maintenance behaviors and “climbing/digging” as manifestation of neglecting behavior.

**Table 1.**
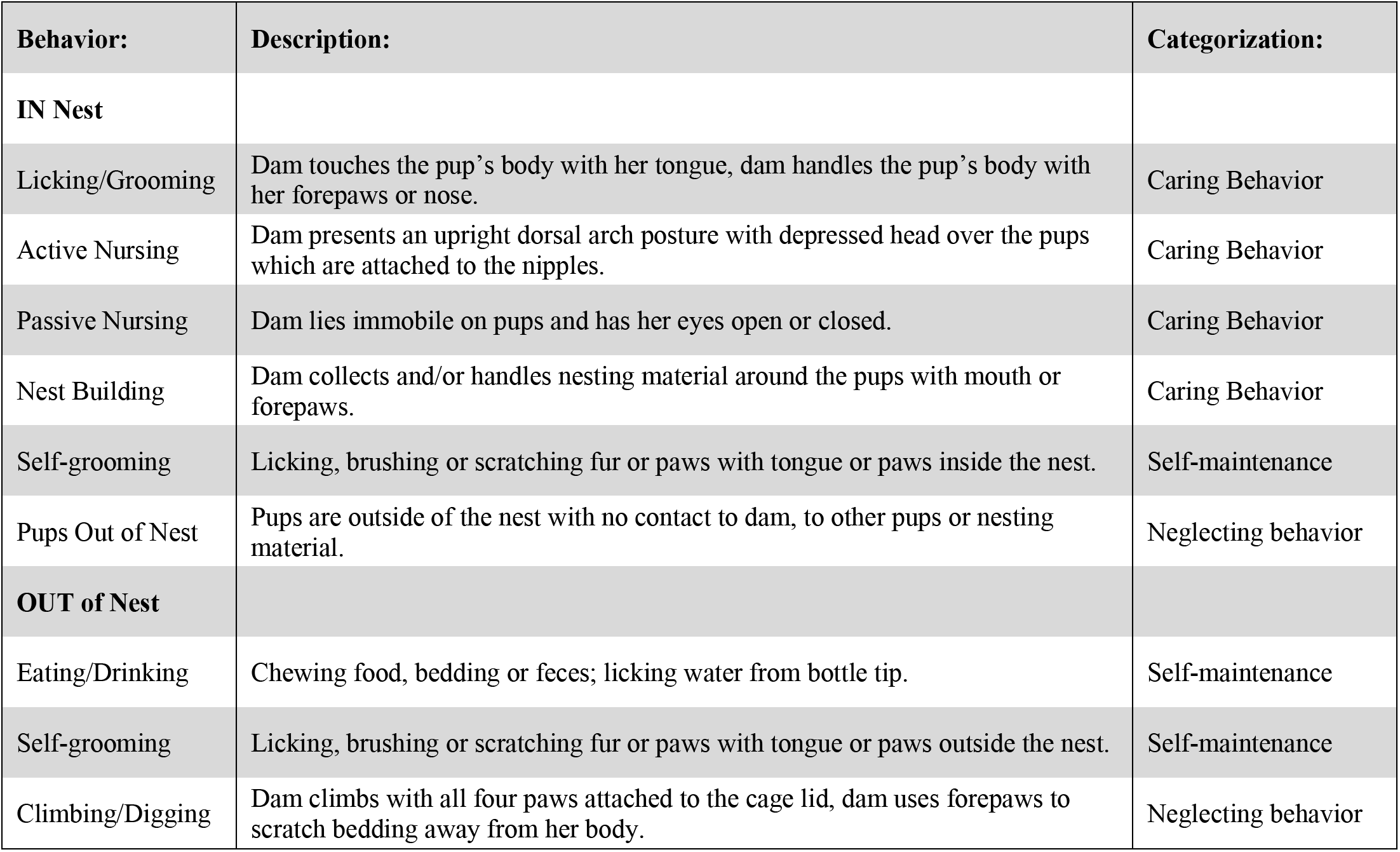
Ethogram used for the assessment of maternal care behavior.

### Pup retrieval test.^36,37^

On PD 7, we briefly removed the dam from the home cage and disturbed the nesting material, distributing it throughout the cage. We then placed two pups in the corners away from the nest end of the home cage. Then, we reintroduced the dam and measured the latency to sniff a pup, retrieve each of the pups, start nest building and crouch over pups. We terminated the test if a dam did not complete the task within 15 minutes, resulting in a latency of 900 seconds for any behaviors not yet observed.

### Spontaneous somatic withdrawal behavior

We tested mice 24 hours after drug cessation on PD 22. We first habituated mice to the testing room for 1 hour, and scored behavior in real time during 5 minute time bins for a 15 minute observation period. Spontaneous somatic withdrawal signs were scored using a modified rating scale.^38,39^ Counted signs included escape jumps and wet dog shakes. Escape jumps numbering 2 to 4 were assigned 1 point, 5 to 9 were assigned 2 points, and 10 or more were assigned 3 points. Wet dog shakes numbering 1 to 2 were assigned 2 points, and 3 or more were assigned 4 points. All other presence signs were assigned 2 points and consisted of 10 distinct behaviors, including persistent trembling, abnormal postures, abnormal gait, paw tremors, teeth chattering, penile erection/genital grooming, excessive eye blinking, ptosis (orbital tightening), swallowing movements, and diarrhea. Counted and presence scores were summed from all 3 of the 5 minute time bins to compute the global withdrawal score.

### Elevated plus maze

This maze is a reliable, canonical test for anxiety-like behavior in rodents.^40^ We habituated mice to the testing room for 1 hour and introduced them to the maze by placing them in the central area, facing one of the open arms, for a 5 minute trial. The time spent in the open and closed arms was recorded using computer tracking software (TopScan CleverSys, Reston, VA).

### Sucrose splash test

The splash test is a pharmacologically validated behavioral assay demonstrated to parallel other affective-like behavioral assays.^41^ We habituated mice to an empty glass cylinder, before spraying their dorsal coat surface 3 times with a 10% sucrose solution. We recorded time spent grooming for 5 minutes.

### Two-tone auditory discrimination

We trained mice daily in two 60 minute sessions (morning and afternoon). Training consisted of four phases: waterspout **habituation** (6 days), behavioral **shaping** (4 days), **single-tone training** (22 days), and **two-tone operant task** (20 days). To motivate task acquisition, mice were mildly water deprived during training days, and received water *ad libitum* on weekends during testing. While mice can readily adapt to water deprivation,^42^ our protocol employed minimal stress. To **habituate** mice to the water spout, water was made available during 10 second trials. We randomized inter-trial intervals (ITI) between 30 and 300 seconds, detecting licks using the Psibox lickspout.^43^ Then, we **shaped** mice to associate a tone with licking and water reward delivery. The target tone was presented for 1 second at 0 dB SPL. The response window began at tone onset and ended after a 3 second post-stimulus silence, with a randomized ITI of 5 to 9 seconds. A conditioning probability was utilized, where 20% of trials were rewarded with 0.5 seconds of water, whether or not the animal responded. To prevent impulsive licking, the subsequent trial was delayed until mice abstained from licking for 5 seconds. During **single-tone training** conditions, an early window was introduced for 0.5 seconds before the start of each trial; if mice responded during this window, a timeout of 20 seconds was added to the ITI. During this phase, the response window was shortened to 2 seconds and the 20% conditioning probability was removed. The parameters for the **two-tone operant task** matched those of single-tone training. We tested mice on 3 distinct target/non-target tone pairs (11000:5000 Hz; 9000:15000 Hz; 7000:13000 Hz) with target to non-target (GO/NOGO) ratios (50/50, 20/80, and 80/20) for 10, 5, and 5 days respectively. Mice received a 20 second timeout if they responded to the non-target tone during the response window. Mice were assigned to respond on a 50/50 target to non-target task to establish basic discrimination differences between groups. Next, we held the target ratio at 20/80 to determine if discrimination can be increased by making rewarding trials less frequent. Finally, we held the target ratio at 80/20 to determine if frequent target trials would decrease discrimination due to the inability to refrain from responding to infrequent non-target trials (i.e., increased false alarms).

### Statistical Analyses

Statistical tests were conducted using Prism 8 (GraphPad) and sample size was determined using G*Power software suite (Heinrich-Heine, Universität Düsseldorf). If no statistically significant sex difference or exposure/sex interaction was apparent, animals were grouped and analyzed according to exposure conditions, per NIH recommendations for studying sex differences. Parametric tests were used when appropriate assumptions were met, otherwise we used nonparametric tests. Cohen’s *d* or η^2^ were used for calculating effect size. All experimenters were blind to treatment conditions throughout data collection, scoring, and analysis.

## Results

### Maternal Care Behavior

We first examined whether there were differences in maternal care. We compared maternal care behavioral observations between fentanyl-exposed and saccharine-exposed (vehicle control) dams (Fig. 1C). Table 1 lists the behaviors used for this assay. Fentanyl-exposed (*n* = 8) and control (*n* = 8) dams had no interaction between behaviors and drug exposure (*F*_(8, 126)_ = 0.63, *p* = 0.74) nor a main effect of drug exposure (*F*_(1, 126)_ = 0.69, *p* = 0.40). These data suggest that fentanyl exposure does not influence maternal care behavior from postpartum to postnatal day 7.

**Figure 1.**
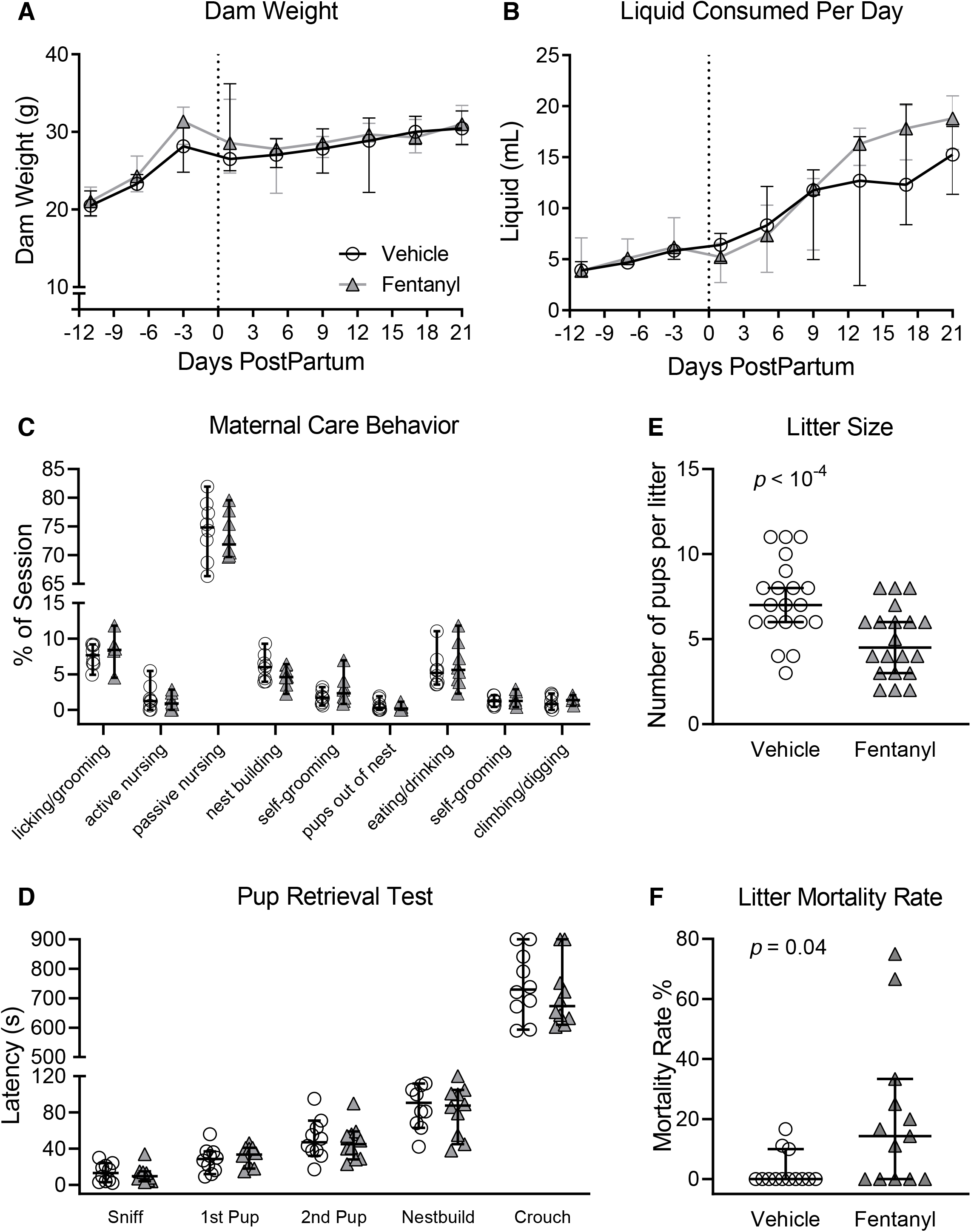
Fentanyl exposure throughout pregnancy and weaning does not influence dam weight (**A**) or daily liquid intake (**B**). **C**: Fentanyl exposure did not adversely affect maternal care behaviors during the critical period from PD 1 to 7. **D**: Fentanyl-exposed dams displayed latencies to pup retrieval on PD 8 that were indistinguishable from controls. Dams exposed to fentanyl during pregnancy had smaller litters (**E**), and a higher litter mortality rate, compared to controls (**F**). Data depict medians and 95% confidence intervals.

To further asses maternal behavior, we used the pup retrieval test, and compared latencies to sniff a pup, retrieve each of two pups, start nest building, and crouch over pups (Fig. 1D). Fentanyl-exposed (*n* = 10) and control (*n* = 10) dams had no interaction between latencies and drug exposure (*F*_(4, 90)_ = 0.35, *p* = 0.83) nor main effect of drug exposure (*F*_(1, 90)_ = 0.64, *p* = 0.42). Together, these data suggest that dams exposed to fentanyl during pregnancy have indistinguishable maternal care behaviors, suggesting that differences in maternal care do not confound the interpretation of the results of fentanyl exposure.

### Dam Weight and Liquid Consumption

Fentanyl exposure does not influence dams’ body weight across pregnancy to weaning (Fig. 1A). We compared total body weight between fentanyl-exposed (*n* = 10) and control (*n* = 10) dams throughout pregnancy until their litters were weaned. There was no interaction between daily weight and drug exposure (*F*_(8, 144)_ = 1.17, *p* = 0.32) or main effect (*F*_(1, 18)_ = 0.38, *p* = 0.55) of fentanyl exposure on the dam weight.

Fentanyl exposure does not influence daily liquid consumption of dams across pregnancy to weaning (Fig. 1B). There was an interaction between daily liquid consumption and drug exposure (*F*_(8, 144)_ = 2.31, *p* = 0.02), however, no difference (*p* > 0.05, Bonferroni) between treatment groups across days. Together, these data suggest that fentanyl exposure during pregnancy does not influence the daily body weight or liquid consumption of dams.

### Litter Size and Mortality Rate

Dams exposed to fentanyl throughout pregnancy until weaning yielded smaller litter sizes (Fig. 1E). Fentanyl-exposed dams (median = 4 pups, 95% CI = 3.89 to 5.380, *n* = 20) had fewer live pups per litter (*p* = 10^−3^, Mann-Whitney, U = 88.50) relative to controls (median = 7 pups, 95% CI = 6.21 to 8.38, *n* = 20), with a large effect size (*d* = 1.12).

Fentanyl exposure during pregnancy increases the mortality rate of litters (Fig. 1F), calculated as a percentage of pups lost from birth until weaning. Litters born to fentanyl-exposed dams (median = 14.29, 95% CI = 5.02 to 35.29, *n* = 13) had a higher mortality rate (*p* = 0.01, Mann-Whitney, U = 43.00) compared to controls (median = 0.00, 95% CI = −0.54 to 6.35, *n* = 13), with a large effect size (*d* = 0.94). These data indicate that dams exposed to fentanyl during pregnancy have smaller litters and have a higher litter mortality rate compared to controls.

### Spontaneous Withdrawal Signs

To test the prediction that cessation of perinatal fentanyl exposure induces somatic withdrawal behavior, we compared global scores of spontaneous withdrawal signs between perinatal fentanyl exposure and control mice, 24 hours after weaning (PD 22, Fig. 2, Table 2). There was an interaction between sex and exposure (*F*_(1, 38)_ = 8.72, *p* < 0.01), with a main effect of sex (*F*_(1, 38)_ = 12.23, *p* < 0.01), and exposure (*F*_(1, 38)_ = 90.42, *p* < 10^−4^), and a large effect size (η^2^ = 0.65). Bonferroni’s post-hoc suggests that exposed male mice (mean = 3.40, 95% CI = 2.43 to 4.36, *n* = 10) exhibited higher withdrawal scores (*p* < 10^−4^) compared to controls (mean = 0.18, 95% CI = −0.08 to 0.45, *n* = 10). Exposed female mice (mean = 2.09, 95% CI = 1.21 to 2.96, *n* = 11) exhibited higher withdrawal scores (*p* = 2 × 10^−4^) compared to controls (mean = 0.18, 95% CI = −0.08 to 0.45, *n* = 11). Additionally, fentanyl exposed male mice exhibited higher withdrawal scores (*p* = 3 × 10^−4^) compared to fentanyl exposed female mice. These data suggest that both male and female mice exposed to fentanyl exhibit spontaneous withdrawal 24 hours after weaning.

**Figure 2.**
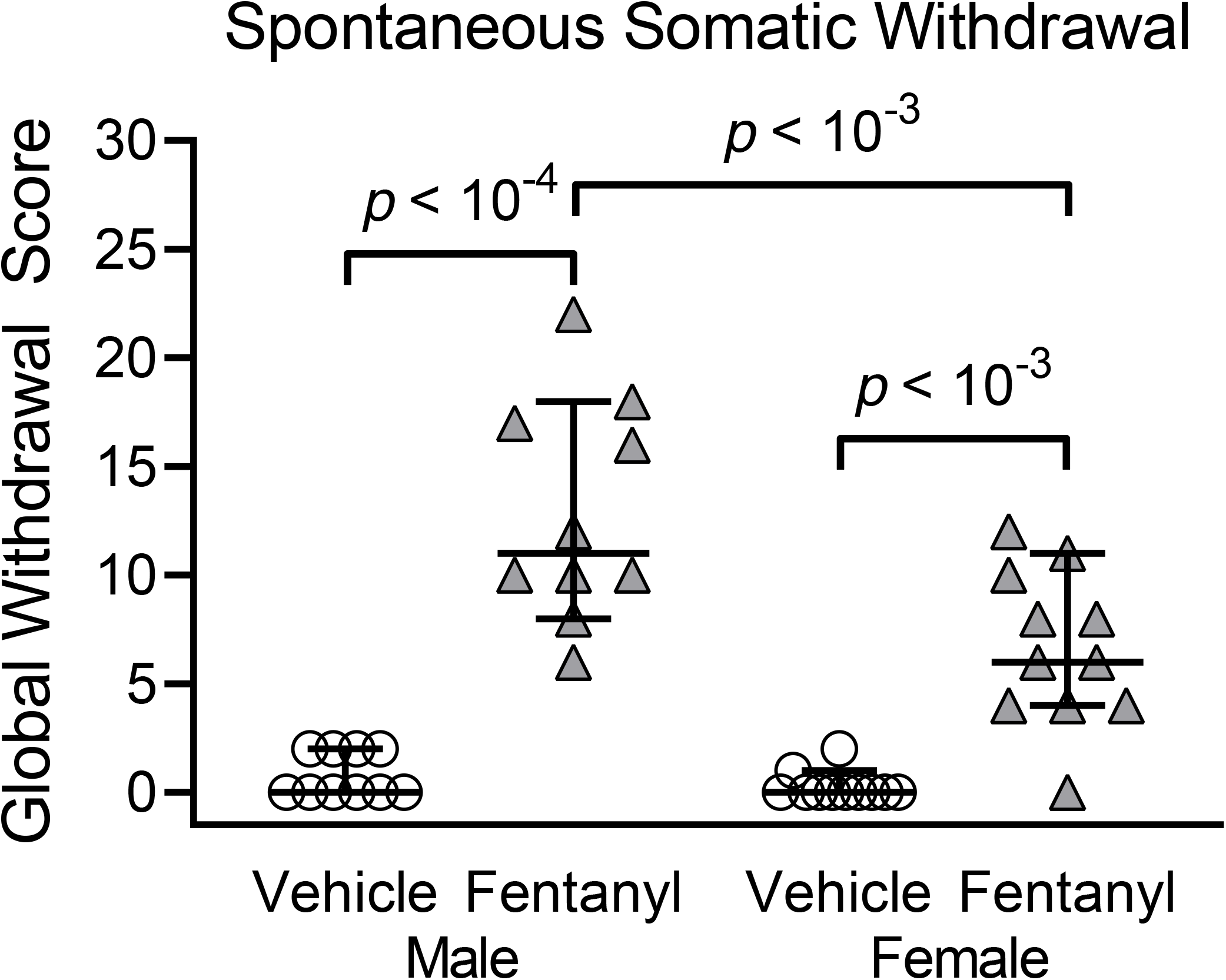
Perinatal fentanyl exposure induces spontaneous somatic withdrawal behavior 24 hours after drug cessation. Exposed mice have higher global withdrawal scores, compared with vehicle controls. Data depict medians and 95% confidence intervals.

**Table 2.**
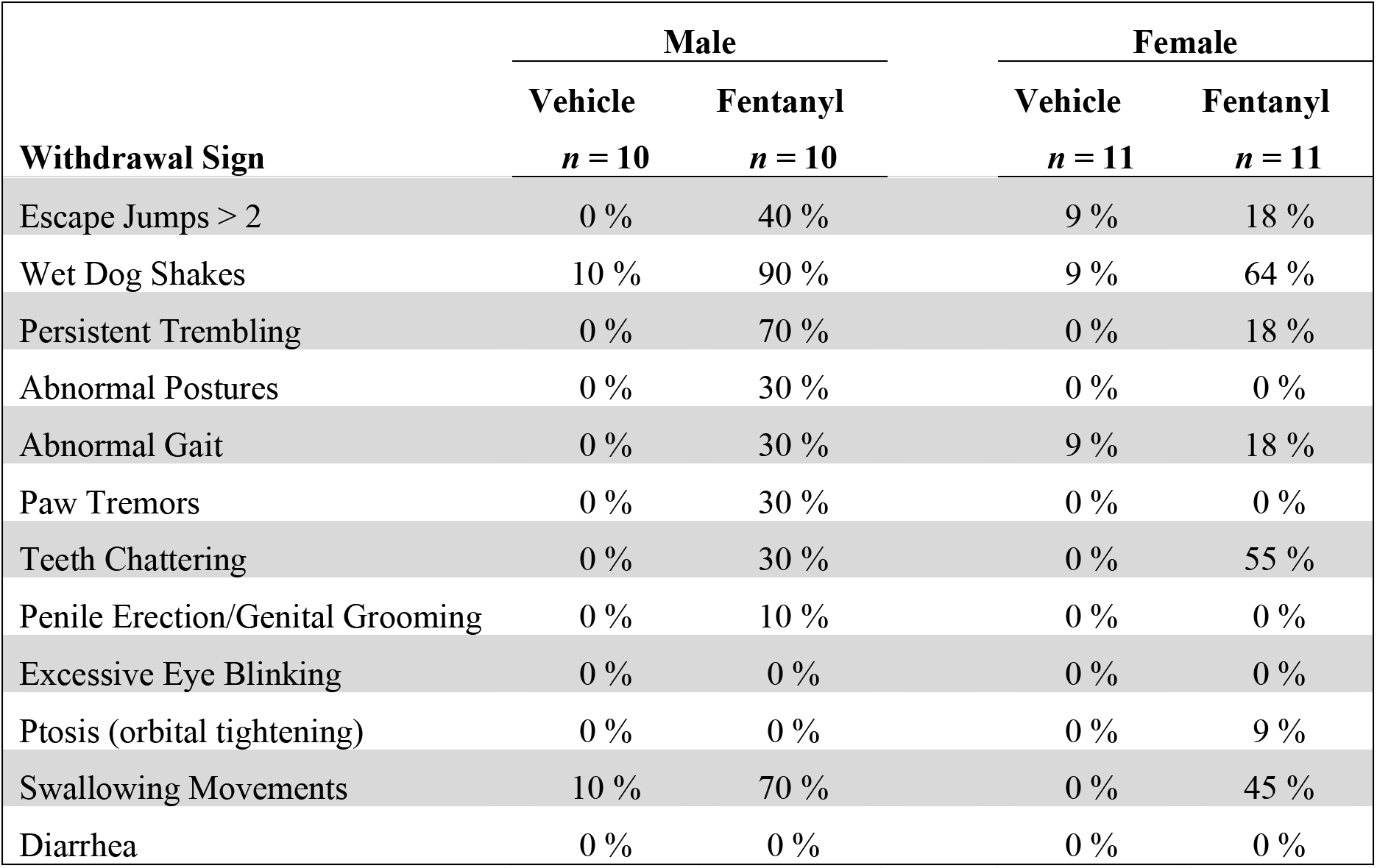
Percentage of mice exhibiting a particular spontaneous somatic withdrawal sign during a 15 minute observation period, 24 hours post drug cessation.

### Litter Weights

To test the prediction that perinatal fentanyl exposure affects body weight across development, we compared the weights at weaning (PD 21), early adolescence (PD 35), and adulthood (PD 55), exposed and control mice.

Figure 3A-C depicts the weight, in grams, of perinatal fentanyl exposure or control mice across development. At weaning (Fig. 3A), exposed male mice weighed less than controls. There was an interaction between sex and drug exposure (*F*_(1, 44)_ = 18.83,*p* < 10^−4^). Tukey’s post hoc comparisons indicate that exposed male mice (mean = 7.39, 95% CI = 7.04 to 7.75; *n* = 12) weighed less (*p* < 10^−4^) than controls (mean = 8.64, 95% CI = 8.27 to 9.01, *n* = 12), with a medium effect size (η^2^ = 0.09). There was no difference (*p* = 0.99) in weight between perinatal fentanyl exposure female mice (mean = 6.39, 95% CI = 6.18 to 6.60, *n* = 12) and controls (mean = 6.43, 95% CI = 6.17 to 6.70, *n* = 12).

By early adolescence (Fig. 3B), both perinatal fentanyl exposed male and female mice weighed more than their respective sex controls. There was no interaction between sex and drug exposure (*F*_(1, 44)_ = 1.27, *p* = 0.27). Bonferroni’s post-hoc suggests that exposed male mice (mean = 19.78, 95% CI = 19.06 to 20.51, *n* = 12) weighed more (*p* = 0.03) than controls (mean = 17.95, 95% CI = 16.87 to 19.03, *n* = 12), with a large effect size (η^2^ = 0.25). Exposed female mice (mean = 17.51, 95% CI = 16.67 to 18.35, *n* = 12) weighed more (*p* = 3 × 10^−4^) than controls (mean = 14.67, 95% CI = 13.44 to 15.89, *n* = 12), with a large effect size (η^2^ = 0.25).

**Figure 3.**
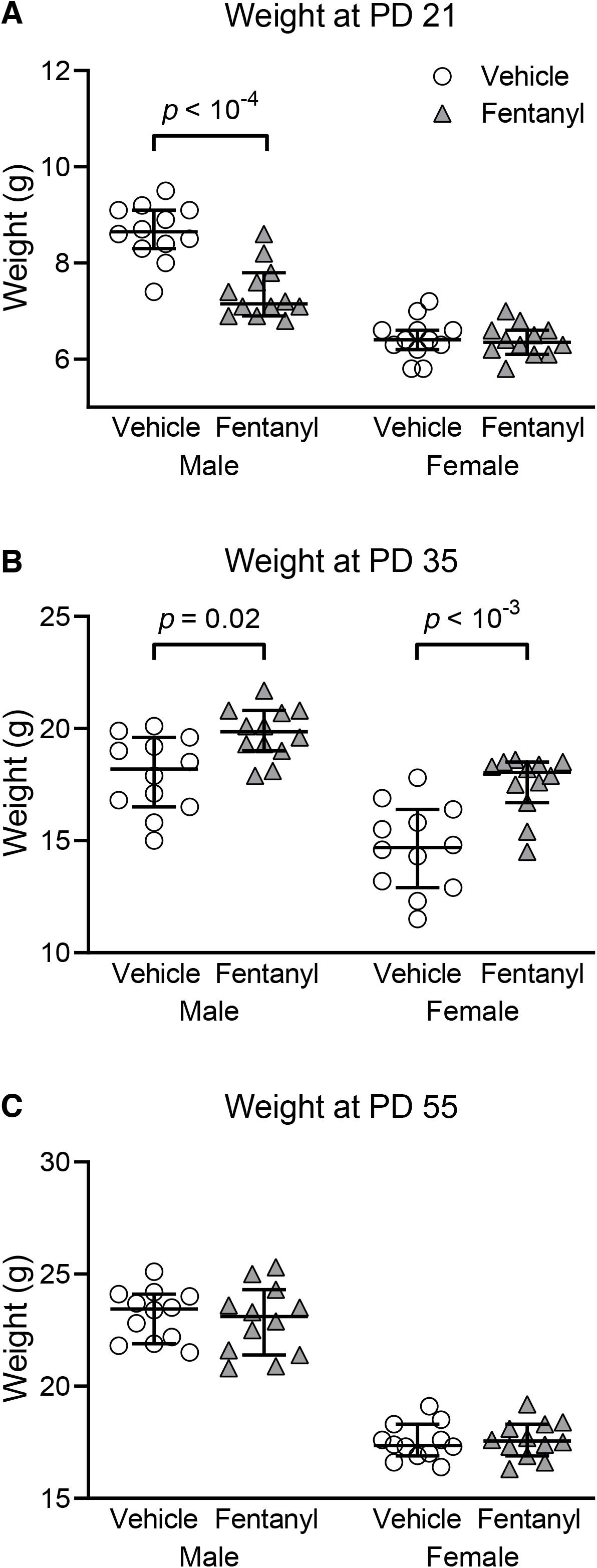
Perinatal fentanyl exposure influences weight across development. **A**: Exposed male mice weigh less than controls at weaning (PD 21). **B**: By adolescence (PD 35), both male and female mice that were exposed to fentanyl weigh more than controls. **C**: This weight difference is no longer present once these mice reach early adulthood (PD 55). Data depict medians and 95% confidence intervals.

By early adulthood (Fig. 3C), there were no weight differences between perinatal fentanyl exposed and control mice (*p* > 0.05). Collectively, these data suggest that male mice exposed perinatally to fentanyl weigh less than controls at weaning. By adolescence, both male and female mice exposed to fentanyl weigh more than controls. This weight difference is no longer present once these mice reach early adulthood.

### Affective-like Behavior

Adolescent children exposed to perinatal opioids exhibit anxiety and aggression.^44^ Therefore, we tested the prediction that perinatal fentanyl exposure influences affective-like behaviors in adolescent mice. We compared the time exposed and control mice spent in the open and closed arms of an elevated plus maze. Exposed male mice displayed more anxiety-like behaviors, compared to controls. When comparing the time spent in the open arms of the maze (Fig. 4A), there was an interaction between sex and exposure (*F*_(1, 62)_ = 4.17, *p* = 0.04), with a main effect of sex (*F*_(1, 62)_ = 10.19, *p* = 0.002). Bonferroni’s post hoc comparisons indicate that exposed male mice (mean = 34.75 sec, 95% CI = 23.20 to 46.29, *n* = 9) spent less time in the open arms (*p* = 0.01) than controls (mean = 50.81 sec, 95% CI = 50.81 to 90.39, *n* = 12), with a small effect size (η^2^ = 0.05). We observed no difference (*p* > 0.99) between exposed female mice (mean = 61.57 sec, 95% CI = 51.50 to 71.63, *n* = 23) and controls (mean = 69.42 sec, 95% CI = 56.94 to 81.90, *n* = 23).

**Figure 4.**
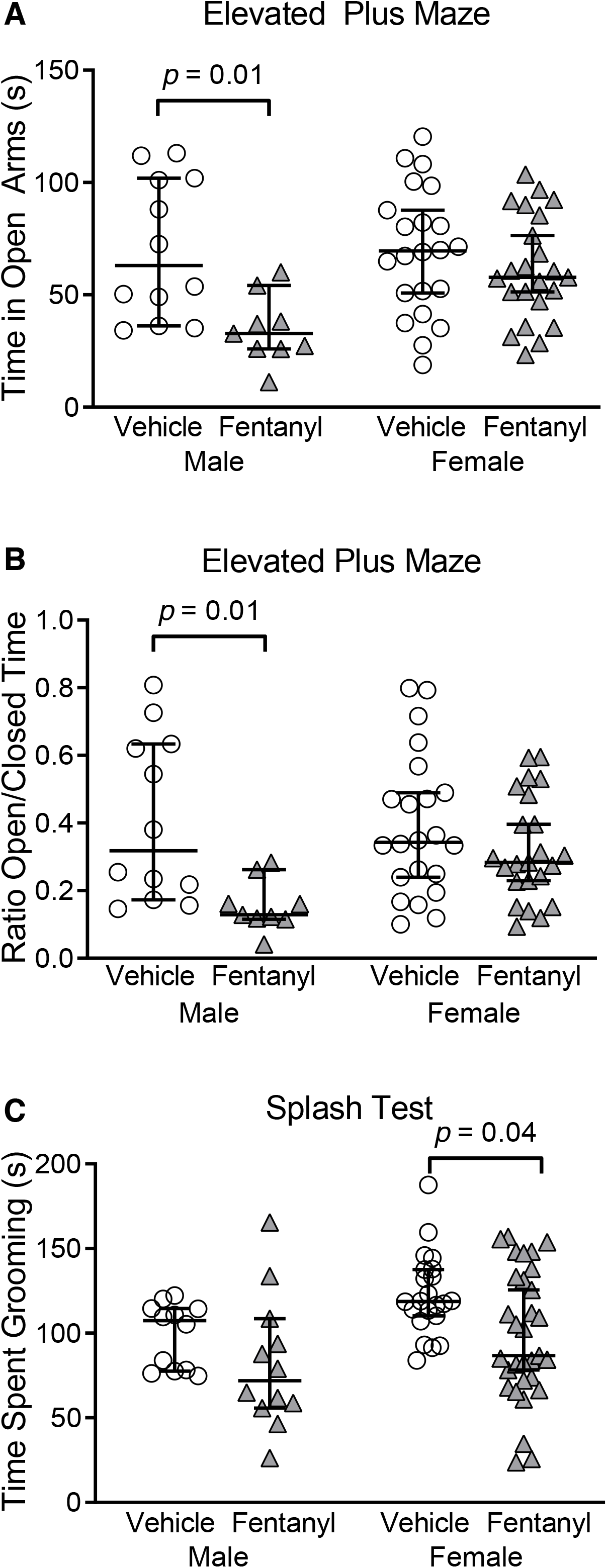
Perinatal fentanyl exposure results in aberrant affective**-**like behaviors in adolescent mice. **A/B**: Adolescent male mice exposed to fentanyl perinatally exhibit increased anxiety-like behavior, assayed with the elevated plus maze. Exposed male mice spend less time in the open arms of the maze, compared to controls. **C**: Adolescent female mice exposed to fentanyl perinatally exhibit decreased grooming behavior, as assayed with the sucrose splash test. Data depict medians and 95% confidence intervals.

We also assessed the difference in the ratio of time spent in the open/closed arms of the elevated plus maze between groups (Fig. 4B). There was no interaction between sex and exposure (*F*_(1, 62)_ = 3.45, *p* = 0.06), however, there was a sex difference (*F*_(1, 62)_ = 10.56, *p* = 0.001). Post hoc comparisons indicate that exposed male mice (mean = 0.15, 95% CI = 0.09 to 0.21, *n* = 9) had a lower ratio (*p* = 0.01), compared to controls (mean = 0.40, 95% CI = 0.25 to 0.56, *n* = 12), with a medium effect size (η^2^ = 0.13). Likewise, there was no difference (*p* = 0.60) in the ratio between exposed female mice (mean = 0.32, 95% CI = 0.25 to 038, *n* = 23) and controls (mean = 0.39, 95% CI = 0.29 to 0.48, *n* = 22). These data suggest that adolescent male mice exposed perinatally to fentanyl exhibit increased anxiety-like behavior.

To assess affective-like behavior, we compared the time exposed and control mice spent grooming during a sucrose splash test (Fig. 4C). Exposed female mice spent less time grooming, compared to controls. There was no interaction between sex and exposure (*F*_(1, 73)_ = 0.19, *p* = 0.66), however, there was a sex difference (*F*_(1, 73)_ = 6.24, *p* = 0.01). Post hoc comparisons indicate that exposed female mice (mean = 98.44 sec, 95% CI = 84.3 to 112.6, *n* = 31) spent less time grooming (*p* = 0.04), compared to controls (mean = 122.70 sec, 95% CI = 112.0 to 133.4, *n* = 22), with a small effect size (η^2^ = 0.07). There was no difference (*p* = 0.57) between exposed male mice (mean = 81.83 sec, 95% CI = 56.96 to 106.7, *n* = 12) and controls (mean = 99.01 sec, 95% CI = 86.91 to 111.1, *n* = 12). Together, these data suggest that adolescent mice exposed perinatally to fentanyl exhibit aberrant affective-like behaviors.

### Auditory Discrimination

Prenatal exposure to opioids impairs inhibitory control in young children.^45^ This is associated with lower performance on perceptual measures of visual, tactile, and auditory tests.^46^ Here, we tested the prediction that perinatal fentanyl exposure will have impaired auditory discrimination and inhibitory control (Fig. 5). Mice were tested on a target to non-target ratio of 20/80 and 80/20, which we predicted would enhance and diminish sensitivity, respectively.^19^ We compared average hits (correctly licked when the target tone was presented), correct rejections (refrained from licking when the non-target tone was presented), total responses (the sum of average hits and false alarms), and *d*’ sensitivity index (a ratio of the hit rate to false alarm rate) between groups. We found that perinatal fentanyl exposure permanently impaired auditory discrimination and task engagement in adult mice.

**Figure 5.**
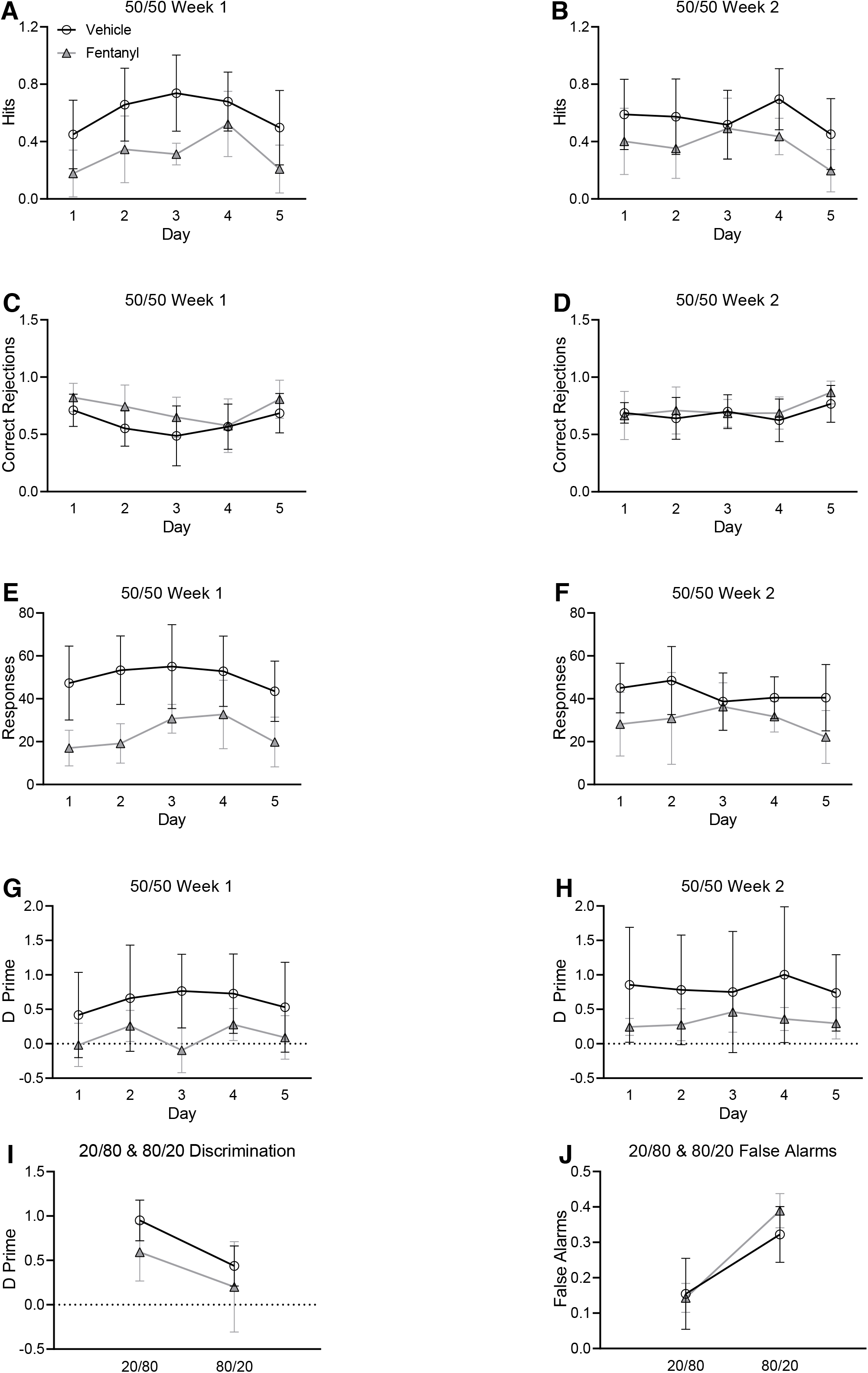
Perinatal fentanyl exposure impairs auditory discrimination and task engagement in adult mice. **A/B**: Exposed mice make fewer correct licks when the target tone is presented. **C/D**: There is no difference between groups in refraining from licking when the non-target tone is presented. **E/F**: perinatal fentanyl exposure mice have fewer responses than vehicle controls. **G/H**: Mice exposed to perinatal fentanyl show lower discrimination sensitivity index, compared to vehicle controls. **I**: All mice have significantly higher discrimination measurements during 20/80 sessions than 80/20 sessions, and there are no differences between groups. **J**: All mice have higher false alarms during 80/20 sessions than 20/80 sessions, and show no difference between groups. Data depict medians and 95% confidence intervals.

Across weeks 1 (Fig. 5A) and 2 (Fig. 5B) of 50/50 training, mice with perinatal fentanyl exposure exhibited fewer correct licks when the target tone was presented, compared to controls. Exposed mice (week 1: mean = 0.31, 95% CI = 0.14 to 0.48; week 2: mean = 0.37, 95% CI = 0.23 to 0.51; *n* = 9) had lower average hits (week 1: *F*_(1, 59)_ = 914.82, *p* = 0.0004; week 2: *F*_(1, 59)_ = 6.79, *p* = 0.01) than controls (week 1: mean = 0.60, 95% CI = 0.45 to 0.75; week 2: mean = 0.56, 95% CI = 0.45 to 0.67; *n* = 9), with a large effect size (week 1: η^2^ = 0.61; week 2: η^2^ = 0.52).

Average correct rejections across weeks 1 (Fig. 5C) and 2 (Fig. 5D) were indistinguishable between groups. Exposed mice (week 1: mean = 0.71, 95% CI = 0.58 to 0.84; week 2: mean = 0.72, 95% CI = .62 to 0.82; *n* = 9) had similar correct rejections (week 1: *F*_(1, 59)_ = 3.99, *p* = 0.05; week 2: *F*_(1, 59)_ = 0.53, *p* = 0.47) as controls (week 1: mean = 0.60, 95% CI = 0.48 to 0.71; week 2: mean = 0.68, 95% CI = 0.61 to 0.75; *n* = 9).

On average, total responses across weeks 1 (Fig. 5E) and 2 (Fig. 5F) were lower in perinatal fentanyl exposure mice compared to controls. Mice perinatally exposed to fentanyl (week 1: mean = 23.87, 95% CI = 14.89 to 32.85; week 2: mean = 29.83, 95% CI = 23.38 to 36.29; *n* = 9) made fewer responses (week 1: *F*_(1, 59)_ = 31.11, *p* = 10^−4^; week 2: *F*_(1, 59)_ = 7.42, *p* = 0.009) than controls (week 1: mean = 50.40, 95% CI = 44.42 to 56.38; week 2: mean = 42.63, 95% CI = 37.63 to 47.63; *n* = 9), with a large effect size (week 1: η^2^ = 0.85; week 2: η^2^ = 0.70).

*d*’ sensitivities measured across weeks 1 (Fig. 5G) and 2 (Fig. 5H) were lower in perinatal fentanyl exposure mice. Across both weeks 1 and 2, exposed mice (week 1: mean = 0.10, 95% CI = −0.10 to 0.30; week 2: mean = 0.32, 95% CI = 0.22 to 0.43; *n* = 9) had decreased sensitivity (week 1: *F*_(1, 59)_ = 9.5, *p* = 0.004; week 2: *F*_(1, 59)_ = 3.73, *p* = 0.02) relative to controls (week 1: mean = 0.62, 95% CI = 0.44 to 0.79; week 2: mean = 0.82, 95% CI = 0.69 to 0.96; *n* = 9), with a large effect size (week 1: η^2^ = 0.78; week 2: η^2^ = 0.89). No interactions were observed across the two weeks of 50/50 sessions (*p* > 0.05).

We also analyzed *d*’ sensitivity measures across the 20/80 to 80/20 (target/non-target) ratios in mice performing the two-tone operant task (Fig. 5I). *d*’ measures were higher in all mice during 20/80 sessions compared to 80/20 (*F*_(1, 97)_ = 12.26, *p* = 0.0007, η^2^ = 0.29). This effect was driven by an increase in false alarms (Fig. 5J, *F*_(1, 97)_ = 30.07, *p* = 10^−4^) but not in hits (*F*_(1, 97)_ = 1.82, *p* = 0.18). During 20/80 sessions, perinatal fentanyl exposure mice made fewer responses, compared to controls (*F*_(1, 59)_ = 4.84, *p* = 0.03). We found no difference (*p* > 0.05) between exposed and control mice in *d*’ sensitivity index nor in false alarm rate during 20/80 or 80/20 sessions.

Taken together, these data suggest that, when the target ratio was held at 50/50, perinatal fentanyl exposure mice made fewer correct responses, fewer overall responses, and exhibited impaired discrimination abilities compared to controls. When we manipulated the frequency of target tones, during 20/80 sessions, exposed mice exhibited fewer responses. However, during 80/20 sessions, it was more difficult for the mice to inhibit licking behavior when the non-target tone was presented. No difference between groups was observed in 80/20 sessions. These data suggest that perinatal fentanyl exposure results in enduring deficits to sensory perception in adult mice.

## Discussion

### Opioid Exposure

Animal models of neonatal opioid withdrawal vary greatly in terms of treatment protocols, species, strain, type of opioid, route of administration, drug dose/concentration, and resulting behavioral alterations. Here, we describe a preclinical model of perinatal fentanyl exposure. Although overall overdose deaths might be slightly decreasing, those from fentanyl continue to rise.^48^ Therefore, it is important to develop and validate preclinical models of perinatal fentanyl exposure.

We chose to administer fentanyl in the drinking water of pregnant dams to better recapitulate the human condition of intermittent opioid use. Exposure to fentanyl through ingestion is not uncommon, as it is responsible for more than half of fentanyl-related overdose deaths.^49^ Furthermore, we wanted to avoid chronic stress involved with repeated injections and handling in mice,^50^ as this might influence maternal care behavior of the dam and later behaviors in offspring.^51^ We also wanted to avoid continuous administration with pumps and pellets, as these do not mimic intermittent use, and they require surgery. Thus, the intermittent nature of dams self-administering fentanyl better models human scenarios.

We administered to the dams fentanyl throughout gestation, and continued this administration until the pups were weaned at PD21. The major neurodevelopmental events occurring during the first 3 postnatal weeks in mice are similar to those occurring during the third trimester of human pregnancy.^3^ Thus, our fentanyl administration mimics human exposure to fentanyl throughout pregnancy.

### Opioid withdrawal

Perhaps the most dramatic and distressing consequence of prenatal opioid exposure in humans is the withdrawal behavior displayed by neonates, collectively referred to as neonatal opioid withdrawal syndrome (NOWS). Commonly observed symptoms include irritability, high-pitched crying, tremors, hypertonicity, vomiting, diarrhea, and tachypnea.^52^

We reasoned that a valid animal model of NOWS should result in a corresponding complement of signs of withdrawal. To our knowledge, few studies assess withdrawal in experimental animals exposed to opioids during the perinatal period,^53^ and none to fentanyl. Here, we demonstrate that perinatal fentanyl exposure results in a large increase in spontaneous somatic withdrawal scores, consistent with the animals exhibiting a NOWS-like phenotype. This is striking considering these withdrawal signs typically require an opioid antagonist, such as naloxone, to precipitate withdrawal.^38^

### Decreased litter size and increased morbidity rate

Opioid use during pregnancy increases the risk of premature births, stillbirths, and sudden infant death.^6^ Rat dams exposed to morphine before and during pregnancy display hormonal imbalances and irregular estrous cycles that are associated with increased litter morbidity.^20,21^ Consistent with these findings, we demonstrate that dams exposed to fentanyl during pregnancy have fewer live pups per litter and a higher morbidity rate.

### Lower birth weights

Human babies exposed to opioids during pregnancy also have lower birth weights,^8^ a finding recapitulated in rats.^20^ Here we show that perinatal fentanyl exposure is associated with lower weights in male mice at weaning, and *higher* weights in adolescents of both sexes. By early adulthood, weights were similar to those of controls. The transient weight increase in adolescence might be specific to fentanyl exposure, the method of exposure we used, or other, not yet known factors.

### Lasting affective deficits

Perinatal opioid exposure in humans results in complications that continue through to adolescence, including increased risk of developing attention deficit hyperactivity disorder,^54^ autism,^55^ autonomic dysregulation,^56^ and poor performance on standardized high school testing.^57^ Children ages 6 to 13 that were exposed to prenatal opioids exhibit affective behavioral deficits.^44^ Analogous outcomes are present in rodent models of early opioid exposure with offspring displaying hyperactivity,^58^ cognitive deficits,^28^ depressive-and anxiety-like behavior.^11,12^ In our model, perinatal fentanyl exposure resulted in aberrant affective behaviors persisting into adolescence, evidenced by their performance on the splash test and the elevated plus maze.

### Lasting sensory deficits

Complications do not end when opioid-exposed babies leave the intensive care unit. Neonates born with NOWS display disruptions in the development of somatosensory networks,^59^ and develop lasting visual,^60^ motor,^61^ and cognitive deficits.^62^ Similarly, rats exposed to neonatal morphine at a time equivalent to the third trimester of gestation, have lasting sensory aberrations, including deficits in nociception, as well as diminished morphine and stress-induced analgesia.^63^ Prenatal morphine exposure inhibits dendritic arborization of cortical pyramidal neurons involved in sensory perception, suggesting long-lasting deficits in sensory processing.^17^ Here, we demonstrate that perinatal fentanyl exposure mice have impaired auditory discrimination and lower levels of task engagement in auditory tasks. Surprisingly, perinatal fentanyl exposure did not appear to impact the ability to refrain responding on non-target trials when there was a prepotency to do so.

These deficits might reflect altered frontal-striatal circuits important for task performance and/or abnormal auditory processing. Impaired auditory processing cannot solely account for the observed differences in engagement because our auditory target and non-target stimuli were loud and highly discriminable, and both groups were impacted by altered ratio schedules in that discrimination improved during 20/80 sessions and worsened during 80/20 sessions. Further, in humans, prenatal opioid exposure is not associated with abnormal auditory evoked brainstem responses, at least in infancy, suggesting that at least lower level auditory processing is intact and that the observed behavioral deficits might originate from altered central auditory processing.^64^ Toddlers with opioid exposure fail to show trial to trial improvements on the 3-box working memory task, suggesting impaired learning mechanisms.^45^ If learning deficits are at play in our study, they appear to persist even with extended testing (20 sessions); discrimination ability of exposed mice did not differ over days in week 2 during 50/50 testing, and discrimination was still impaired during the last week of testing when ratio manipulations were performed.

### Conclusions

We describe a preclinical, rodent model of perinatal fentanyl exposure that recapitulates key aspects of the human condition. This model may allow mechanistic studies of the lasting consequences of perinatal exposure to this potent and widely-used opioid, to enable the development of approaches, and to prevent or ameliorate these consequences.

## Acknowledgements

Olivia Uddin and Lace M. Riggs (Program in Neuroscience, University of Maryland School of Medicine) critically reviewed content and provided revision.

## Author contribution

AK, MKL, MRR, POK were responsible for the study concept and design. JBA, ATB, MEF, SST, and CAd conducted the experiments. JBA, ATB, and MEF performed data analysis. JBA, ATB, MRR, and AK drafted the manuscript. DE, NAF, POK, MRR, and AK provided critical revision of the manuscript for content. All authors critically reviewed content and approved the final version for publication.

## References

1. Vivolo-Kantor AM, Seth P, Gladden RM, et al. Vital Signs: Trends in Emergency Department Visits for Suspected Opioid Overdoses – United States, July 2016-September 2017. MMWR Morb Mortal Wkly Rep. 2018;67(9):279–285. doi:10.15585/mmwr.mm6709e1

2. Florence CS, Zhou C, Luo F, Xu L. The Economic Burden of Prescription Opioid Overdose, Abuse, and Dependence in the United States, 2013. Med Care. 2016;54(10):901–906. doi:10.1097/MLR.0000000000000625

3. Ross EJ, Graham DL, Money KM, Stanwood GD. Developmental consequences of fetal exposure to drugs: what we know and what we still must learn. Neuropsychopharmacol. 2015;40(1):61–87. doi:10.1038/npp.2014.147

4. Mars SG, Rosenblum D, Ciccarone D. Illicit fentanyls in the opioid street market: desired or imposed? Addict Abingdon Engl. December 2018. doi:10.1111/add.14474

5. Spencer MR, Warner M, Bastian BA, Trinidad JP, Hedegaard H. Drug Overdose Deaths Involving Fentanyl, 2011-2016. Natl Cent Health Stat Natl Vital Stat Syst. 2019;68(3):1–19.

6. Whiteman VE, Salemi JL, Mogos MF, Cain MA, Aliyu MH, Salihu HM. Maternal opioid drug use during pregnancy and its impact on perinatal morbidity, mortality, and the costs of medical care in the United States. J Pregnancy. 2014;2014:906723. doi:10.1155/2014/906723

7. Holbrook A, Kaltenbach K. Gender and NAS: does sex matter? Drug Alcohol Depend. 2010;112(1-2):156–159. doi:10.1016/j.drugalcdep.2010.05.015

8. Kandall SR, Albin S, Lowinson J, Berle B, Eidelman AI, Gartner LM. Differential effects of maternal heroin and methadone use on birthweight. Pediatrics. 1976;58(5):681–685.

9. Ornoy A, Michailevskaya V, Lukashov I, Bar-Hamburger R, Harel S. The developmental outcome of children born to heroin-dependent mothers, raised at home or adopted. Child Abuse Negl. 1996;20(5):385–396.

10. Castellano C, Ammassari-Teule M. Prenatal exposure to morphine in mice: enhanced responsiveness to morphine and stress. Pharmacol Biochem Behav. 1984;21(1):103–108. doi:10.1016/0091-3057(84)90138-2

11. Byrnes EM. Transgenerational consequences of adolescent morphine exposure in female rats: effects on anxiety-like behaviors and morphine sensitization in adult offspring. Psychopharmacology (Berl). 2005;182(4):537–544. doi:10.1007/s00213-005-0122-4

12. Klausz B, Pintér O, Sobor M, et al. Changes in adaptability following perinatal morphine exposure in juvenile and adult rats. Eur J Pharmacol. 2011;654(2):166–172. doi:10.1016/j.ejphar.2010.11.025

13. Lu R, Liu X, Long H, Ma L. Effects of prenatal cocaine and heroin exposure on neuronal dendrite morphogenesis and spatial recognition memory in mice. Neurosci Lett. 2012;522(2): 128–133. doi:10.1016/j.neulet.2012.06.023

14. Vathy IU, Etgen AM, Rabii J, Barfield RJ. Effects of prenatal exposure to morphine sulfate on reproductive function of female rats. Pharmacol Biochem Behav. 1983;19(5):777–780. doi:10.1016/0091-3057(83)90079-5

15. Gagin R, Cohen E, Shavit Y. Prenatal exposure to morphine alters analgesic responses and preference for sweet solutions in adult rats. Pharmacol Biochem Behav. 1996;55(4):629–634.

16. Laborie C, Dutriez-Casteloot I, Montel V, Dickès-Coopman A, Lesage J, Vieau D. Prenatal morphine exposure affects sympathoadrenal axis activity and serotonin metabolism in adult male rats both under basal conditions and after an ether inhalation stress. Neurosci Lett. 2005;381(3):211–216. doi:10.1016/j.neulet.2005.01.083

17. Mei B, Niu L, Cao B, Huang D, Zhou Y. Prenatal morphine exposure alters the layer II/III pyramidal neurons morphology in lateral secondary visual cortex of juvenile rats. Synap N Y N. 2009;63(12): 1154–1161. doi:10.1002/syn.20694

18. Ahmadalipour A, Sadeghzadeh J, Vafaei AA, Bandegi AR, Mohammadkhani R, Rashidy-Pour A. Effects of environmental enrichment on behavioral deficits and alterations in hippocampal BDNF induced by prenatal exposure to morphine in juvenile rats. Neuroscience. 2015;305:372–383. doi:10.1016/j.neuroscience.2015.08.015

19. Verde ME, MacMillan NA, Rotello CM. Measures of sensitivity based on a single hit rate and false alarm rate: the accuracy, precision, and robustness of d’, Az, and A’. Percept Psychophys. 2006;68(4):643–654.

20. Siddiqui A, Haq S, Shaharyar S, Haider SG. Morphine induces reproductive changes in female rats and their male offspring. Reprod Toxicol. 1995;9(2):143–151. doi:10.1016/0890-6238(94)00064-6

21. Siddiqui A, Haq S, Shah BH. Perinatal exposure to morphine disrupts brain norepinephrine, ovarian cyclicity, and sexual receptivity in rats. Pharmacol Biochem Behav. 1997;58(1):243–248. doi:10.1016/s0091-3057(97)00012-9

22. Chiou L-C, Yeh G-C, Fan S-H, How C-H, Chuang K-C, Tao P-L. Prenatal morphine exposure decreases analgesia but not K+ channel activation. Neuroreport. 2003;14(2):239–242. doi:10.1097/00001756-200302100-00016

23. Hou Y, Tan Y, Belcheva MM, Clark AL, Zahm DS, Coscia CJ. Differential effects of gestational buprenorphine, naloxone, and methadone on mesolimbic mu opioid and ORL1 receptor G protein coupling. Brain Res Dev Brain Res. 2004;151(1-2):149–157. doi:10.1016/j.devbrainres.2004.05.002

24. Huleihel R, Yanai J. Disruption of the development of cholinergic-induced translocation/activation of PKC isoforms after prenatal heroin exposure. Brain Res Bull. 2006;69(2): 174–181. doi :10.1016/j.brainresbull.2005.11.023

25. Schrott LM, Franklin L ‘Tonya M, Serrano PA. Prenatal opiate exposure impairs radial arm maze performance and reduces levels of BDNF precursor following training. Brain Res. 2008;1198:132–140. doi:10.1016/j.brainres.2008.01.020

26. Chiang Y-C, Hung T-W, Lee CW-S, Yan J-Y, Ho I-K. Enhancement of tolerance development to morphine in rats prenatally exposed to morphine, methadone, and buprenorphine. J Biomed Sci. 2010;17:46. doi:10.1186/1423-0127-17-46

27. Wu C-C, Hung C-J, Shen C-H, et al. Prenatal buprenorphine exposure decreases neurogenesis in rats. Toxicol Lett. 2014;225(1):92–101. doi:10.1016/j.toxlet.2013.12.001

28. Chen H-H, Chiang Y-C, Yuan ZF, et al. Buprenorphine, methadone, and morphine treatment during pregnancy: behavioral effects on the offspring in rats. Neuropsychiatr Dis Treat. 2015;11:609–618. doi:10.2147/NDT.S70585

29. Devarapalli M, Leonard M, Briyal S, et al. Prenatal Oxycodone Exposure Alters CNS Endothelin Receptor Expression in Neonatal Rats. Drug Res. 2016;66(5):246–250. doi:10.1055/s-0035-1569279

30. Chen VS, Morrison JP, Southwell MF, Foley JF, Bolon B, Elmore SA. Histology Atlas of the Developing Prenatal and Postnatal Mouse Central Nervous System, with Emphasis on Prenatal Days E7.5 to E18.5. Toxicol Pathol. 2017;45(6):705–744. doi:10.1177/0192623317728134

31. Frisina RD, Singh A, Bak M, Bozorg S, Seth R, Zhu X. F1 (CBA×C57) mice show superior hearing in old age relative to their parental strains: hybrid vigor or a new animal model for “golden ears”? Neurobiol Aging. 2011;32(9):1716–1724. doi:10.1016/j.neurobiolaging.2009.09.009

32. Wade CL, Schuster DJ, Domingo KM, Kitto KF, Fairbanks CA. Supraspinally-administered agmatine attenuates the development of oral fentanyl self-administration. Eur J Pharmacol. 2008;587(1-3):135–140. doi:10.1016/j.ejphar.2008.04.007

33. Wade CL, Krumenacher P, Kitto KF, Peterson CD, Wilcox GL, Fairbanks CA. Effect of Chronic Pain on Fentanyl Self-Administration in Mice. Taylor B, ed. PLoS ONE. 2013;8(11):e79239. doi:10.1371/journal.pone.0079239

34. Tordoff MG, Bachmanov AA, Reed DR. Forty mouse strain survey of water and sodium intake. Physiol Behav. 2007;91(5):620–631. doi:10.1016/j.physbeh.2007.03.025

35. Murko M, Elek B, Styblo M, Thomas DJ, Francesconi KA. Dose and Diet – Sources of Arsenic Intake in Mouse in Utero Exposure Scenarios. Chem Res Toxicol. 2018;31(2):156–164. doi:10.1021/acs.chemrestox.7b00309

36. Chourbaji S, Hoyer C, Richter SH, et al. Differences in mouse maternal care behavior – is there a genetic impact of the glucocorticoid receptor? PloS One. 2011;6(4):e19218. doi:10.1371/journal.pone.0019218

37. Champagne FA, Curley JP, Keverne EB, Bateson PPG. Natural variations in postpartum maternal care in inbred and outbred mice. Physiol Behav. 2007;91(2-3):325–334. doi:10.1016/j.physbeh.2007.03.014

38. Luster BR, Cogan ES, Schmidt KT, et al. Inhibitory transmission in the bed nucleus of the stria terminalis in male and female mice following morphine withdrawal. Addict Biol. April 2019. doi:10.1111/adb.12748

39. Schulteis G, Heyser CJ, Koob GF. Differential expression of response-disruptive and somatic indices of opiate withdrawal during the initiation and development of opiate dependence. Behav Pharmacol. 1999;10(3):235–242. doi:10.1097/00008877-199905000-00001

40. Pellow S, Chopin P, File SE, Briley M. Validation of open:closed arm entries in an elevated plus-maze as a measure of anxiety in the rat. J Neurosci Methods. 1985;14(3):149–167.

41. Reis-Silva TM, Sandini TM, Calefi AS, et al. Stress resilience evidenced by grooming behaviour and dopamine levels in male mice selected for high and low immobility using the tail suspension test. Eur J Neurosci. 2019;50(6):2942–2954. doi:10.1111/ejn.14409

42. Bekkevold CM, Robertson KL, Reinhard MK, Battles AH, Rowland NE. Dehydration parameters and standards for laboratory mice. J Am Assoc Lab Anim Sci JAALAS. 2013;52(3):233–239.

43. Francis NA, Kanold PO. Automated Operant Conditioning in the Mouse Home Cage. Front Neural Circuits. 2017;11. doi:10.3389/fncir.2017.00010

44. de Cubas MM, Field T. Children of methadone-dependent women: developmental outcomes. Am J Orthopsychiatry. 1993;63(2):266–276.

45. Levine TA, Woodward LJ. Early inhibitory control and working memory abilities of children prenatally exposed to methadone. Early Hum Dev. 2018;116:68–75. doi:10.1016/j.earlhumdev.2017.11.010

46. Wilson GS, McCreary R, Kean J, Baxter JC. The development of preschool children of heroin-addicted mothers: a controlled study. Pediatrics. 1979;63(1):135–141.

47. Wu P-L, Yang Y-N, Suen J-L, Yang Y-CSH, Yang C-H, Yang S-N. Long-Lasting Alterations in Gene Expression of Postsynaptic Density 95 and Inotropic Glutamatergic Receptor Subunit in the Mesocorticolimbic System of Rat Offspring Born to Morphine-Addicted Mothers. BioMed Res Int. 2018;2018:5437092. doi:10.1155/2018/5437092

48. Ahmad F, Escobedo L, Rossen L, Spencer M, Warner M, Sutton P. National Vital Statistics System: Provisional Drug Overdose Death Counts. Centers for Disease Control and Prevention; 2019. https://www.cdc.gov/nchs/nvss/vsrr/drug-overdose-data.htm. Accessed July 30, 2019.

49. O’Donnell JK, Halpin J, Mattson CL, Goldberger BA, Gladden RM. Deaths Involving Fentanyl, Fentanyl Analogs, and U-47700 – 10 States, July-December 2016. MMWR Morb Mortal Wkly Rep. 2017;66(43):1197–1202. doi:10.15585/mmwr.mm6643e1

50. Ryabinin AE, Wang Y-M, Finn DA. Different Levels of Fos Immunoreactivity After Repeated Handling and Injection Stress in Two Inbred Strains of Mice. Pharmacol Biochem Behav. 1999;63(1):143–151. doi:10.1016/S0091-3057(98)00239-1

51. Soares-Cunha C, Coimbra B, Borges S, et al. Mild Prenatal Stress Causes Emotional and Brain Structural Modifications in Rats of Both Sexes. Front Behav Neurosci. 2018;12:129. doi:10.3389/fnbeh.2018.00129

52. Finnegan LP, Connaughton JF, Kron RE, Emich JP. Neonatal abstinence syndrome: assessment and management. Addict Dis. 1975;2(1-2):141–158.

53. Tao PL, Yeh GC, Su CH, Wu YH. Co-administration of dextromethorphan during pregnancy and throughout lactation significantly decreases the adverse effects associated with chronic morphine administration in rat offspring. Life Sci. 2001;69(20):2439–2450. doi:10.1016/s0024-3205(01)01316-9

54. Ornoy A, Segal J, Bar-Hamburger R, Greenbaum C. Developmental outcome of school-age children born to mothers with heroin dependency: importance of environmental factors. Dev Med Child Neurol. 2001;43(10):668–675.

55. Rubenstein E, Young JC, Croen LA, et al. Brief Report: Maternal Opioid Prescription from Preconception Through Pregnancy and the Odds of Autism Spectrum Disorder and Autism Features in Children. J Autism Dev Disord. 2019;49(1):376–382. doi:10.1007/s10803-018-3721-8

56. Paul JA, Logan BA, Krishnan R, et al. Development of auditory event-related potentials in infants prenatally exposed to methadone. Dev Psychobiol. 2014;56(5):1119–1128. doi:10.1002/dev.21160

57. Oei JL, Melhuish E, Uebel H, et al. Neonatal Abstinence Syndrome and High School Performance. Pediatrics. 2017;139(2):e20162651. doi:10.1542/peds.2016-2651

58. Sithisarn T, Bada HS, Charnigo RJ, Legan SJ, Randall DC. Effects of perinatal oxycodone exposure on the cardiovascular response to acute stress in male rats at weaning and in young adulthood. Front Physiol. 2013;4:85. doi:10.3389/fphys.2013.00085

59. Kivistö K, Nevalainen P, Lauronen L, Tupola S, Pihko E, Kivitie-Kallio S. Somatosensory and auditory processing in opioid-exposed newborns with neonatal abstinence syndrome: a magnetoencephalographic approach. J Matern-Fetal Neonatal Med. 2015;28(17):2015–2019. doi:10.3109/14767058.2014.978755

60. Gupta M, Mulvihill AO, Lascaratos G, Fleck BW, George ND. Nystagmus and Reduced Visual Acuity Secondary to Drug Exposure In Utero: Long-Term Follow-up. J Pediatr Ophthalmol Strabismus. 2012;49(1):58–63. doi:10.3928/01913913-20110308-01

61. Bunikowski R, Grimmer I, Heiser A, Metze B, Schäfer A, Obladen M. Neurodevelopmental outcome after prenatal exposure to opiates. Eur J Pediatr. 1998;157(9):724–730.

62. Guo X, Spencer JW, Suess PE, Hickey JE, Better WE, Herning RI. Cognitive brain potential alterations in boys exposed to opiates: In utero and lifestyle comparisons. Addict Behav. 1994;19(4):429–441. doi:10.1016/0306-4603(94)90065-5

63. Zhang GH, Sweitzer SM. Neonatal morphine enhances nociception and decreases analgesia in young rats. Brain Res. 2008;1199:82–90. doi:10.1016/j.brainres.2007.12.043

64. Grimmer I, Bührer C, Aust G, Obladen M. Hearing in newborn infants of opiate-addicted mothers. Eur J Pediatr. 1999;158(8):653–657.

